# Functional selective FPR1 signaling in favor of an activation of the neutrophil superoxide generating NOX2-complex

**DOI:** 10.1101/2020.05.20.106310

**Authors:** Simon Lind, Claes Dahlgren, Rikard Holmdahl, Peter Olofsson, Huamei Forsman

## Abstract

Two formyl peptide receptors (FPR1 and FPR2), abundantly expressed by neutrophils, regulate both pro-inflammatory tissue recruitment of neutrophils and resolution of inflammatory reactions. This dual functionality of the FPRs, opens for a possibility to develop receptor selective therapeutics as mechanism for novel anti-inflammatory treatments. In line with this, high throughput screening studies have identified numerous FPR ligands belonging to different structural classes, but a potent FPR1 agonist with defined biased signaling and functional selectivity has not yet been reported. In this study, we used an FPR1 selective small compound agonist (RE) that represents a chemical entity developed from NOX2 activators identified from our earlier screening studies (WO2012127214). This FPR1 agonist potently activates neutrophils to produce reactive oxygen species (ROS, EC_50_ ~1 nM), whereas it is a weaker chemoattractant than the prototype FPR1 agonist fMLF. At the signaling level, RE has a strong bias towards the PLC-PIP_2_-Ca^2+^ pathway and ERK1/2 activation but away from β-arrestin recruitment and the ability to recruit neutrophils chemotactically. In addition, FPR1 when activated by RE could cross-regulate other receptor-mediated neutrophil functions. In comparison to the peptide agonist fMLF, RE is more resistant to oxidization-induced inactivation by the MPO-H_2_O_2_-halide system. In summary, this study describes as a novel FPR1 agonist displaying a biased signaling and functional selectivity when activating FPR1 in human blood neutrophils. RE could possibly be a useful tool compound not only for further mechanistic studies of the regulatory role of FPR1 in inflammation *in vitro* and *in vivo*, but also for developing FPR1specific drug therapeutics.

## 1. Introduction

Neutrophils express several G protein-coupled receptors (GPCRs) that regulate cell functions and fine-tune inflammatory reactions [1, 2]. Among these receptors, the chemoattractant formyl peptide receptors (FPR1 and FPR2) have gained much interest over the years and they have been extensively studied by researchers both in academia and pharmaceutical industry [3-5]. FPR1 and FPR2 are strongly associated with the progression as well as the resolution of inflammatory reactions initiated by microbial infections and/or aseptic tissue injuries [6, 7]. The receptors recognize not only microbial pathogen associated molecular patterns and host derived danger signals in the form of formylated peptides, but also numerous non-formylated peptides/proteins/lipopeptides and other molecules such as small compounds and peptidomimetics [3, 8, 9]. FPR1 and FPR2 exhibit a large overall amino acid sequence similarity with a high degree of identity in the cytosolic parts and a lower in the extracellular domains [4]. This suggests that the two receptors differ more when it comes to ligand binding than in the intracellular signals transmitted. Nevertheless, many agonists cross activate the two receptors although there are a few reported that are highly specific for one or the other of the two receptors [9]. The down-stream signals generated by agonists of FPRs regulate neutrophil directional migration (chemotaxis), mobilization of adhesion molecules to the cell surface, secretion of inflammatory mediators including proteolytically active proteases. Another feature of FPR agonists is also the activation of the electron transporting NADPH-oxidase complex type 2 (the NOX2 complex) with the capacity to generate superoxide anions (O_2_^−^) that form also other reactive oxygen species (ROS) [10].

Activation of neutrophils is essential for defence against microbes and for clearance of harmful tissue debris, but also to limit further neutrophil recruitment and facilitate tissue repair. Thus, these dual functions needs to be tightly controlled through the different phases of inflammation. The effect of ROS shows a similar type of complex role at different phases and at different types of inflammation. ROS released in high quantities from neutrophils is generally regarded as driving acute inflammation. However, it is also clear that low ROS could exaggerate inflammation, effects that are more likely to operate in the resolution phases [11-13]. Hence, in light of this complex and so far not completely understood role of FPR and ROS regulation, our accumulated research proposes a regulatory role of ROS produced by the NADPH-oxidase in many cellular processes [14, 15]. Patients, as well as experimental animals, with chronic granulomatous disease (CGD), lacking the ability to generate ROS, suffer not only from severe microbial infections, but also from a variety of inflammatory complications indicative of important functions of ROS in the mechanisms that control inflammation [16-18]. The importance of ROS in the regulation of inflammation also gains support from our earlier studies in which we through positional cloning of a disease linked genetic polymorphism, have identified *Ncf1* (encoding for the p47^phox^ subunit of the NADPH-oxidase complex) as a disease-associated gene [19] and the molecular basis being linked to a compromised ROS production [20]. Similarly, polymorphism of *Ncf1* plays a role in human autoimmune diseases [21, 22], and in animal models, it has been shown to be of importance for disease severity of arthritis, psoriasis, colitis, and lupus, reviewed in [14, 23]. It is apparent from both pharmacological and genetic deletion studies, that FPRs have multiple roles in diseases conditions associated with a dysregulated inflammation. Mice deficient in individual Fprs show not only an increased susceptibility to microbial infections but also a delayed tissue repair [7, 24, 25]. In addition, a recent study has elegantly demonstrated that activation of FPRs improves cardiac function in a post myocardial infarction model [26], suggesting an anti-inflammatory/pro-resolving role of FPR agonists.

The introduction of the biased GPCR signaling concept rapidly became the starting point not only for more detailed characterization of known GPCR agonists but also for the search for new biased GPCR agonists that could be used to develop drug candidates [27-29]. The concept of biased signaling and functional selectivity give at hand that related to the agonist that binds, an activated GPCR can be stabilized in a conformation that allows or blocks one of the multiple signaling cascades that trigger receptor down-stream functional activities, and the concept has been shown to be valid also for FPR2. This is clearly illustrated by the down-stream signaling by FPR2 specific agonistic lipopeptides/pepducins, peptidomimetics as well as by formylated peptides generated by virulent *S. aureus* bacteria [30-32]. These biased FPR2 agonists are potent in triggering a rise in intracellular calcium ([Ca^2+^]_i_) and release of superoxide through the NADPH-oxidase but they lack ability to recruit β-arrestin and induce chemotaxis [30-32]. It is reasonable to assume, that also the agonist occupied FPR1 can be stabilized in a conformation that opens for one signaling pathway downstream of the receptor but not for another. This assumption gains support from the study showing that selective formylpeptide analogues can discriminate between different biological responses, being able to trigger chemotaxis but not to activate the superoxide generating neutrophil NADPH-oxidase [33].

In attempt to identify ROS activators, we have earlier screened libraries of small compounds and identified a number of hits belonging to different structural classes (patent WO2012127214; [34]). Among these hits, one lead compound has been further developed as a novel compound class of structures (represented by the FPR agonist RE-04-001/RE) that activates differentiated neutrophil-like HL60 cells with an activation profile that is very similar to well-characterized FPR agonists. This observation promoted us to hypothesize that RE could be an FPR agonist and a detailed characterization of the compound was performed in this study; the data revealed it to be an FPR1 agonist inducing a functional selective neutrophil response. This functional selectivity was closely linked to a biased signaling feature in favor of ERK1/2 phosphorylation and rise of intracellular Ca^2+^ together with an inability to recruit β-arrestin. The biased agonistic profile of RE, being a potent activator of ROS production, suggests that RE could serve as a valuable representative novel compound for further mechanistic studies designed to dissect the contribution of different FPR1-mediated functions in inflammation associated diseases as well as potential therapeutic agent.

## 2. Material and Methods

### 2.1 Ethics Statement

This study, conducted at the Sahlgrenska Academy in Sweden, includes peripheral blood and from buffy coats obtained from the blood bank at Sahlgrenska University Hospital, Gothenburg, Sweden. According to the Swedish legislation section code 4§ 3p SFS 2003:460 (Lag om etikprövning av forskning som avser människor), no ethical approval was needed since the blood samples were provided anonymously and could not be traced back to a specific donor.

### 2.2 Chemicals and reagents

The compound RE-04-001 (RE), with structure related to the quinolones that were described in the patent application WO 2012/127214 and reported earlier in screening studies [34]. For intellectual property reasons, the chemical structure of RE is not disclosed, but to make it possible to reproduce the data presented herein, the compound will be provided to other researchers under a standard material transfer agreement (contact).

Dextran T500 was obtained from Pharmacocosmos (Holbaek, Denmark), Ficoll-Paque was from GE Healthcare Bio-Science AB (Uppsala, Sweden), and Fura-2-AM was from Life Technologies Europe (Stockholm, Sweden). RPMI 1640 culture medium without phenol red was purchased from PAA Laboratories GmbH (Pasching, Austria). Isoluminol, N-formyl-Met-Leu-Phe (fMLF), cetyltrimethylammonium bromide (CTAB), o-Phenylenediamine (OPD), EGTA, dimethyl sulfoxide (DMSO), platelet activating factor (PAF), bovine serum albumin (BSA) and Latrunculin A were obtained from Sigma-Aldrich (St. Louis, MO, USA). Horse radish peroxidase (HRP) was purchased from Boehringer-Mannheim (Mannheim, Germany). TNFα and IL8 were from R&D systems (Minneapolis, MN, USA). PAF was from Calbiochem. The FPR2 agonist WKYMVM was synthesized and purified by HPLC by Alta Bioscience (University of Birmingham, Birmingham, UK). The FPR2 specific antagonist PBP10 was synthesized by CASLO Laboratory (Lyngby, Denmark) and the FPR1 specific inhibitor (an inverse agonist) cyclosporin H was kindly provided by Novartis Pharma (Basel, Switzerland). The Gαq inhibitor YM-254890 was purchased from Wako Chemicals (Neuss, Germany). Myeloperoxidase (MPO) and the phenylacetamide compound (S)-2-(4-chlorophenyl)-3,3-dimethyl-N-(5-phenylthiazol-2-yl) butanamide (Cmp58) was obtained from Calbiochem-Merck Millipore (Billerica, MA, USA). Compound 43 was from Tocris Bioscience. The Act-389949 compound [35], synthesized by Ramidus AB (Lund, Sweden) are generous gift from ProNoxis AB (Lund, Sweden). The peptides/receptor antagonists were dissolved in DMSO to a concentration of 10^−2^ M and stored at −80°C until use. Further dilutions were made in Krebs-Ringer phosphate buffer containing glucose (10 mM), Ca^2+^ (1 mM), and Mg^2+^ (1.5 mM) (KRG; pH 7.3). The small compound RE-04-001, was a generous gift from ProNoxis AB (Lund, Sweden). For more information about RE-04-001, please contact the corresponding author Peter Olofsson.

#### 2.2. Isolation of human neutrophils and culture of neutrophil-like HL-60 cells

Neutrophil granulocytes were isolated from peripheral blood or buffy coats obtained from healthy adults [36, 37]. After dextran sedimentation at 1 × *g*, hypotonic lysis of the remaining erythrocytes, and centrifugation on a Ficoll-Paque gradient, the neutrophils were washed and re-suspended (1 × 10^7^/ml) in KRG. The cells were stored on melting ice until used. The purity of the neutrophil preparations was routinely >90%. HL60 cells were cultured under sterile conditions at 37° C in 5% CO_2_ in RPMI 1640 medium supplemented with 10% fetal calf serum (FCS), 2 mM L-glutamine, 1 mM sodium pyruvate, 100 units/mL penicillin and 100 μg/mL streptomycin (RPMI-complete medium). Cells were cultured at a density of 2 × 10^5^ cells/ml in tissue culture flasks (75 cm^2^) and differentiated towards a non-adherent neutrophil-like phenotype by incubation with 1% DMSO for five days. Cells were washed and re-suspended to 10^6^/ml in KRG, stored on ice until use on day five after start of the differentiation.

### 2.3 Calcium mobilization

Neutrophils at a density of 5 × 10^7^ cells/ml in KRG without Ca^2+^ supplemented with 0.1% BSA were loaded with Fura-2-AM (5 μM) for 30 minutes in the dark at room temperature. The cells were then diluted 1:1 in RPMI 1640 culture medium without phenol red and centrifuged at 900 rpm × *g*. Finally, the cells were washed once with KRG and re-suspended in the same buffer to a density of 2 × 10^7^/ml. Calcium measurements were carried out in a PerkinElmer fluorescence spectrophotometer (LC50, Perkin Elmer, Waltham, MA, USA), with excitation wavelengths of 340 nm and 380 nm, an emission wavelength of 509 nm, and slit widths of 5 nm and 10 nm, respectively. The transient rise in intracellular calcium is presented as the ratio of fluorescence intensities (340 nm/380 nm) detected.

### 2.4 Neutrophil NADPH-oxidase activity

Neutrophil superoxide anion production was determined using an isoluminol-enhanced chemiluminescence (CL) system (details are given in [38]). The CL activity was measured in a six-channel Biolumat LB 9505 (Berthold Co, Wildbad, Germany) using disposable 4-ml polypropylene tubes with a 1-ml reaction mixture. Tubes containing isoluminol (2 × 10^−5^ M), HRP (2 units/ml), and neutrophils (10^5^/ml) were equilibrated for five minutes at 37°C, after which 0.1 ml of stimuli was added and the superoxide production, measured as light emission and expressed in Mega counts per minute (Mcpm), was recorded continuously over time.

### 2.5 Treatment of FPR agonists with MPO-H_2_O_2_

Different peptide or small compound FPR agonist was incubated with MPO (1 μg/ml) at ambient temperature for five minutes before the addition of H_2_O_2_ (10 μM final concentration), and incubation was continued for another 10 min at ambient temperature to allow peptide oxidation. The remaining activity of the agonists after MPO-H_2_O_2_-halide oxidation was determined through the potential of agonist to trigger ROS release from neutrophils. The control agonists were incubated at the same concentration in KRG but with no addition of MPO and H_2_O_2_.

### 2.6 Chemotaxis assay

Neutrophil migration was determined by a Boyden chamber technique using 96-well microplate chemotaxis chambers containing polycarbonate filters with 3 μm pores (Chemo-Tx; Neuro Probe Inc., Gaithersburg, MD, USA) according to manufacturer’s instructions. In short, RE-04-001, fMLF or WKYMVM diluted in KRG buffer supplemented with 0.3% BSA, were added to wells in the lower chamber. Cell suspensions (30 μl) containing neutrophils (2 × 10^6^/ml, isolated from peripheral blood) were placed on top of the filter and allowed to migrate for 90 minutes at 37°C. The cell migration to the bottom well was visualized under microscope and for quantitative analysis the content of MPO was assessed in the lysates (cells in lower chamber treated with 2% BSA and 2% CTAB for 60 minutes, at room temperature) by addition of OPD and hydrogen peroxide. Triplicate samples for each stimulus was performed and all values were subtracted by the negative control (spontaneous migration of neutrophils towards buffer). The data is presented as percent migration as compared to the positive control (neutrophils added directly to the bottom chamber, i.e., 100% migration).

### 2.7 β-arrestin 2 recruitment assay

The ability of agonists in promoting FPRs to β-arrestin was evaluated in the PathHunter^®^ eXpress CHO-K1 FPR1 or FPR2 cells from DiscoverX (Fremont, CA, USA) which co-express ProLink tagged FPR1 or FPR2 and an enzyme acceptor tagged β-arrestin so that β-arrestin binding can be measured via enzyme fragment complementation as increased β-galactosidase activity. The assay was performed according to manufacturer’s instructions as previously described [31, 35]. In brief, cells were seeded in tissue culture treated 96-well plates (10^4^ cells/well) and incubated at 37°C, 5% CO_2_ for 20 hours. The cells were then incubated with agonists (90 minutes, 37°C), followed by addition of detection solution and incubated for another 60 minutes at room temperature. The chemiluminescence was measured on a CLARIOstar plate reader (BMG Labtech, Ortenberg, Germany).

### 2.8. Phosphorylation of ERK1/2 determined by electrochemiluminescence

Human neutrophils (2 × 10^6^/ml) were stimulated with fMLF or RE for 2 min followed by rapidly cool down to stop reaction with ice old lysis buffer provided by Meso Scale Diagnostics (MSD, Rockville, Maryland) according to manufacturer’s instructions as described [32]. The lysis was performed on ice for at least 30 min and the supernatant was collected and stored at −80°C before use. Phosphorylation of ERK1/2 (pERK) was measured using the MSD electrochemiluminescence technology with

### 2.9 Data analysis

Data analysis was performed using GraphPad Prism 8.0 (Graphpad Software, San Diego, CA, USA). Curve fitting was performed by non-linear regression using the sigmoidal dose-response equation (variable-slope). Statistical analysis was performed on raw data values using either a repeated measurement one-way ANOVA followed by Dunnett’s multiple comparison post-hoc test or a paired Student’s *t*-test. Statistically significant differences are indicated by \**p* < 0.05, \*\**p* < 0.01

## 3. Results

### 3.1 RE, identified through its ability to activate neutrophil-like HL60 cells, activates also human neutrophils determined as a rise in intracellular Ca^2+^ and FPR1 is the recognizing receptor

A compound library containing drug-like small molecules was used in a high throughput screening study to identify novel NADPH-oxidase activators [34]. The release of superoxide anions from neutrophil-like HL60 cells was determined, and RE was found to activate the NADPH-oxidase and with an activation pattern similar to that of FPR activating compound (Fig 1). The response induced by RE shared a similar activation profile both in magnitude and time course, to that induced by the two high-affinity FPR agonists fMLF (specific for FPR1) and WKYMVM (specific for FPR2) (Fig 1).

**Fig. 1.**
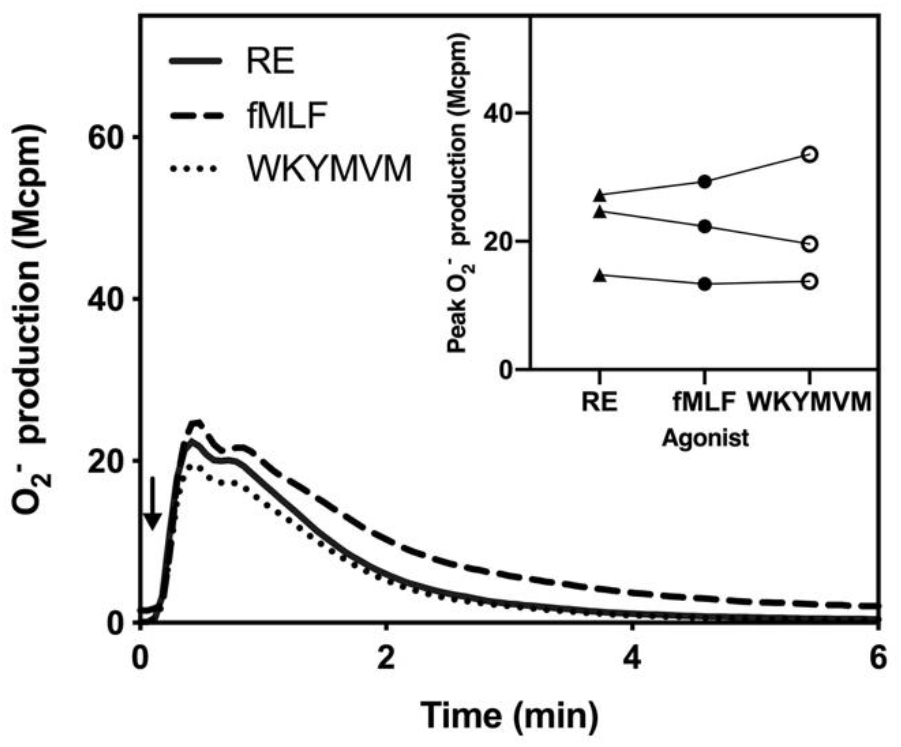
The novel small compound RE-04-001 (denoted as RE in all figures) activates neutrophil-like HL60 cells. The ability of RE to activate DMSO-differentiated neutrophil-like HL60 cells to produce ROS was measured by isoluminol-amplified chemiluminescence technique. The response induced by RE (100 nM, solid line) was compared to that induced by the FPR1 agonist fMLF (100 nM, dashed line) and FPR2 agonist WKYMVM (100 nM, dotted line). Cells were pre-incubated at 37°C for five minutes before agonist stimulation (indicated by arrows) and the NADPH-oxidase mediated superoxide anion (O_2_^−^) production was measure over time. Abscissa, Time (min); ordinate, O_2_^−^ production, arbitrary Mcpm units). **Inset**: comparison of the magnitude (the peak O_2_^−^ production) induced by RE and the two FPR agonists. Each symbol represents an individual experiment.

It is well known that both FPR1 and FPR2 are abundantly expressed by human neutrophils and the receptors recognize numerous structurally unrelated agonists [3, 9]. The similarity in the responses both in kinetics and in magnitude, induced by RE and the two FPR peptide agonists (Fig 1) promoted us to hypothesize that RE may be an FPR agonist that should activate also primary blood neutrophils isolated from healthy donors. One of the very early signaling events down-stream of activated neutrophil FPRs, is a transient increase in the intracellular concentration of free calcium ions ([Ca^2+^]_i_), an event initiated by a G-protein dependent activation of phospholipase C and a release of Ca^2+^ from intracellular storage organelles [39]. Hence, we could show that also RE induced a robust and concentration dependent rise in [Ca^2+^]_i_ in human neutrophils (Fig 2A). In comparison to the prototype FPR peptide agonists fMLF and WKYMVM, RE was the more potent; a full response was obtained already with a 1 nM concentration, and the activity was retained even at concentrations down to 0.1 nM, a concentration unable to trigger a rise in [Ca^2+^]_i_ with the commonly used prototype FPR peptide agonists fMLF or WKYMVM (Fig 2B). Also, in the Ca^2+^ assay system using human neutrophils, RE and the FPR peptide agonists triggered very similar response, further suggesting that RE may interact with FPRs to mediate its biological responses in neutrophils. To determine the involvement of FPRs in the RE induced neutrophil activation, we used two well-known receptor-specific antagonist cyclosporine H (antagonizes primarily FPR1; [40]) and PBP10 (antagonizes primarily FPR2; [41]). The results obtained with these antagonists clearly show that FPR1 was involved in mediating RE induced rise in [Ca^2+^]_i_ (Fig 2C). Based on the lack of inhibition with the FPR2 antagonist, we conclude that FPR2 is not of importance in mediating the RE response (Fig 2C). For comparison, control experiments with prototype peptide agonists for FPR1 and FPR2 are included to show that cyclosporine H selectively inhibits the fMLF-induced response, whereas PBP10 inhibits the WKYMVM response without any effect on the fMLF-induced response (Fig 2C).

**Fig. 2.**
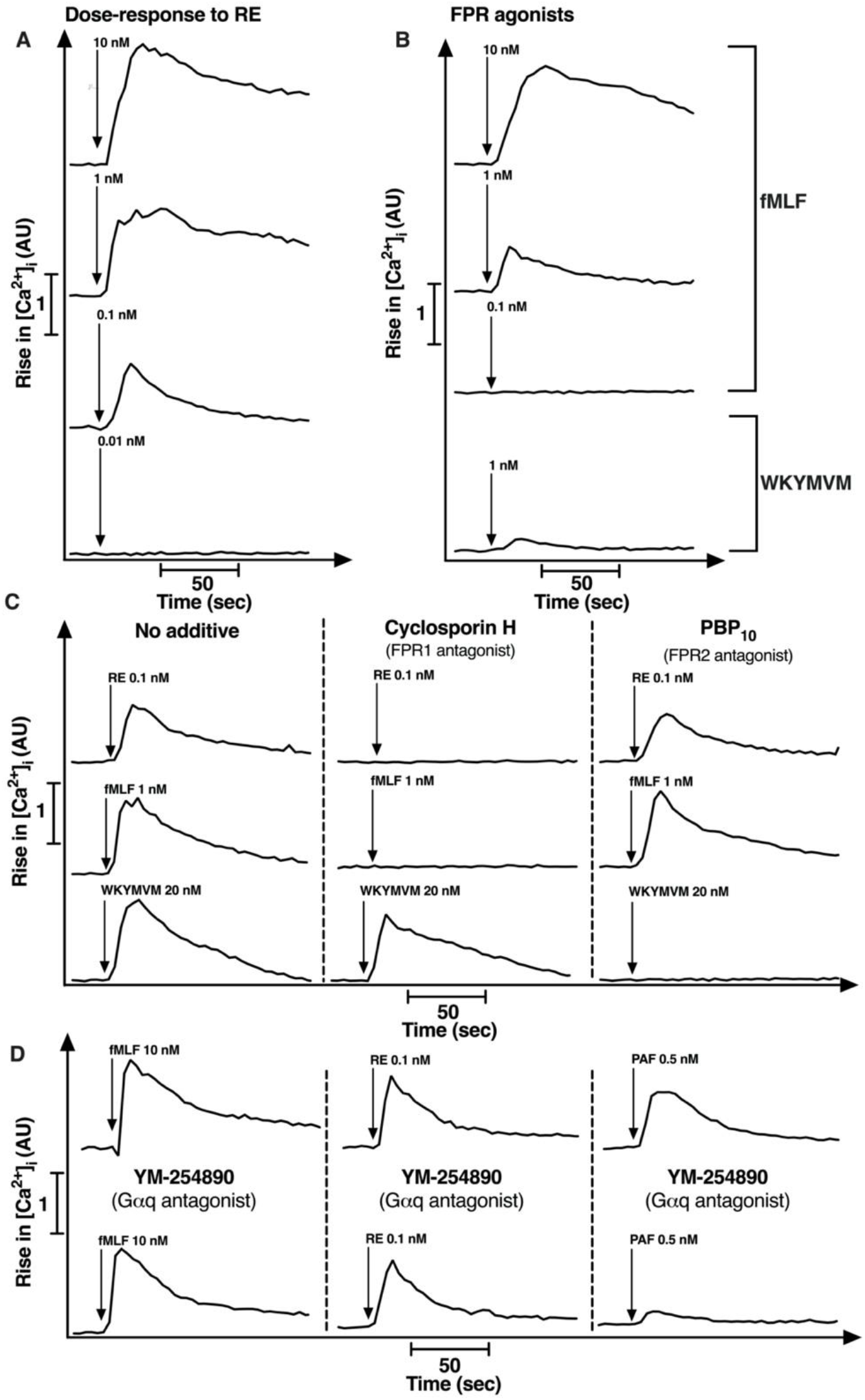
The small compound RE triggers FPR1-mediated intracellular rise of Ca^2+^ independent of Gαq protein activation in human neutrophils. Neutrophils were loaded with Fura-2 and stimulated with different agonists added as indicated by arrows. Abscissa, time of study (sec); ordinate, increase in intracellular Ca^2+^ ([Ca^2+^]_i_) given as the change in the ratio between Fura-2 fluorescence at 340 and 380 nm (AU, arbitrary units). **A-B)** The transient rise of [Ca^2+^]_i_ in neutrophils was induced by different concentrations of RE (10 nM to 0.01 nM), the FPR1 peptide agonist fMLF (10 to 0.1 nM) and the FPR2 agonist WKYMVM (1 nM). **C)** Effect of the FPR1 antagonist cyclosporin H (1 μM, middle panel) or the FPR2 antagonist PBP10 (1 μM, right panel) on the transient rise of [Ca^2+^]_i_ induced by RE (0.1 nM). No antagonist addition before agonist stimulation was used as control (left panel). Agonist addition fMLF (1nM) and WKYMVM (20 nM) were included for comparison. Agonist addition was indicated by arrows. **D)** Effect of YM-254890 (a selective Gαq inhibitor; 200 nM) on the RE response. Neutrophils were left untreated (upper panel) or pre-treated with YM-254890 (lower panel) for five minutes before stimulation with fMLF (10 nM), RE (0.1 nM) or PAF (0.5 nM). The transient rise of intracellular Ca^2+^ was monitored. **A-D)** Representative traces from 3 independent experiments are shown (n = 3).

For many GPCRs, the transient rise in [Ca^2+^]_i_ upon agonist binding is achieved through an activation of a Gαq containing G protein followed by the activation of downstream PLC-PIP_2_-IP_3_ pathway leading to the emptying of intracellular Ca^2+^ stores [42, 43].

One such Gαq-linked neutrophil receptor is the platelet activating factor receptor (PAFR) [44], and accordingly, the PAF-induced rise in [Ca^2+^]_i_ was inhibited by the selective Gαq inhibitor YM-254890 (Fig 2D). In contrast to the PAFR, the FPR-mediated [Ca^2+^]_i_ response does not engage Gαq but the heterodimeric Gβγ subunit derived from a Gαi containing G protein [44]. The fact that the rise in [Ca^2+^]_i_ induced by RE was insensitive to the Gαq selective inhibitor (Fig 2D), is in line with the notion that RE interacts with FPR1 and that the [Ca^2+^]_i_ rise is achieved through a Gαi containing G protein. Taken together, these data clearly show that RE activates human neutrophils manifested as a rise in [Ca^2+^]_i_, and the response is sensitive to an antagonist of FPR1-but not to one for FPR2- or to a Gαq-selective inhibitor.

### 3.2 RE activates neutrophils to release of superoxide anions

To further assess neutrophil activation by RE, we determined the ability of the compound to trigger an assembly of the O_2_^−^ generating NADPH-oxidase in human neutrophils. We show that RE activates neutrophils to release O_2_^−^, and there was a very rapid onset of the response that was then terminated in around 5 min after the initiation, a response pattern very similar to that induced by the two prototype FPR peptide agonists (Fig 3A). The maximal level of O_2_^−^ production induced by RE was of the same magnitude as that induced by 100 nM fMLF, suggesting that RE is a full agonist (Fig 3A). The response induced by RE was concentration dependent with an EC_50_ value in the low nano-molar range (Fig 3B) which is much lower than that for the prototype FPR1 agonist fMLF (≈ 20 nM; Fig 3B). In line with the data obtained with FPR specific antagonists in the [Ca^2+^]_i_ assay system (Fig 2C), the inhibitory profile for RE was the same as that that of fMLF (sensitive to cyclosporine H but not to PBP_10_) but different from WKYMVM (Fig 3C).

**Fig. 3.**
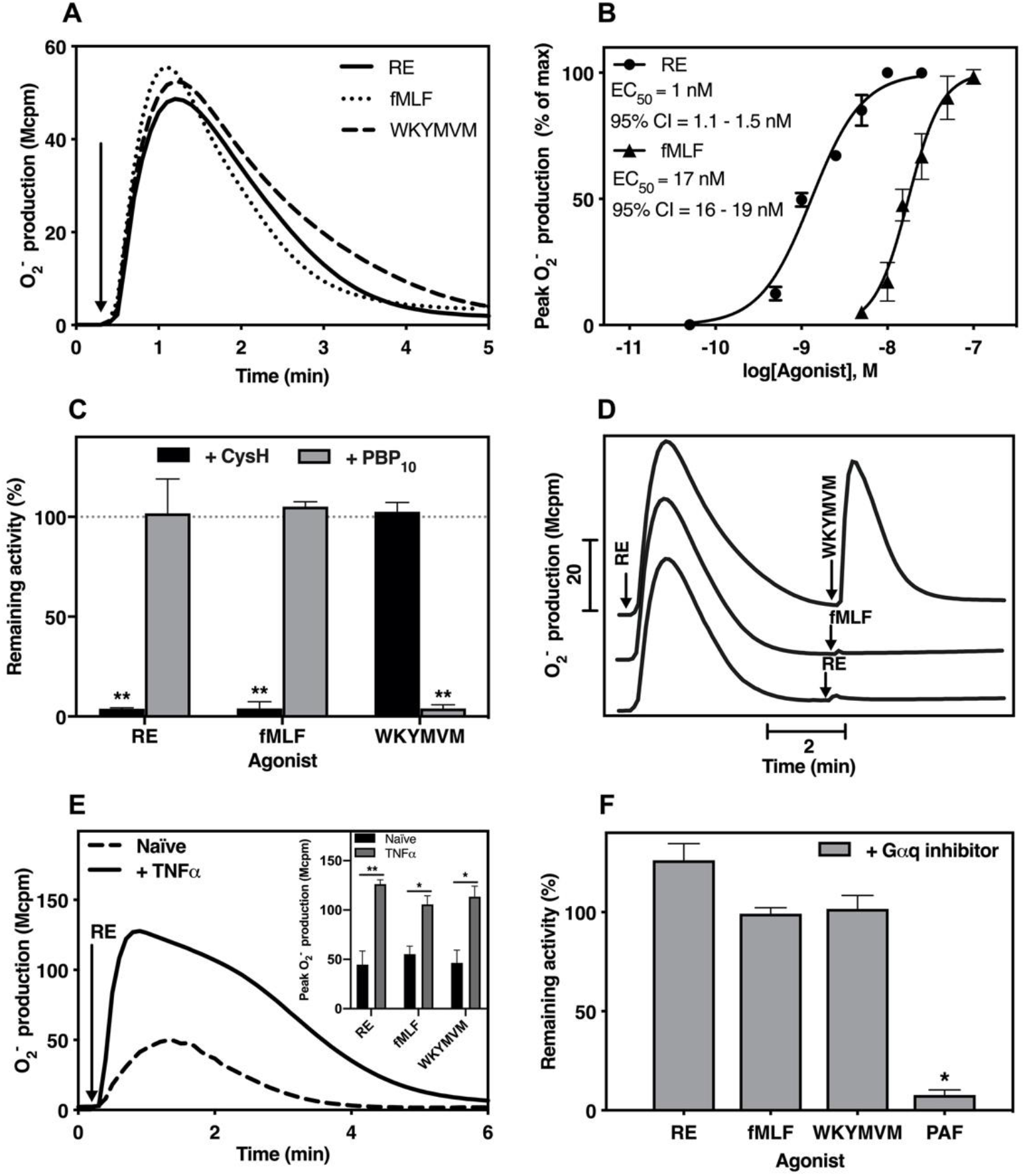
The small compound RE induces FPR1-mediated NADPH-oxidase activation independent of Gαq protein activation from human neutrophils. Neutrophils were pre-incubated at 37°C for five min before stimulation (indicated by arrows) and the NADPH-oxidase mediated O_2_^−^ production was determined. Abscissa, Time (min); ordinate, O_2_^−^ production, arbitrary Mcpm units). **A)** Neutrophils were stimulated with RE (10 nM, solid line), fMLF (100 nM, dotted line) or WKYMVM (100 nM, dashed line). One representative trace out of three independent experiments is shown. **B)** Dose-response of RE and fMLF from 3 independent experiments. The EC_50_-values and 95% confidence interval (CI) were determined based on the peak O_2_^−^ response. **C)** Comparison between the peak O_2_^−^ responses released by neutrophils pretreated without or with either cyclosporin H (1 μM, black bars) or PBP10 (1 μM, grey bars) for five minutes before activation with RE (10 nM), fMLF (100 nM) or WKYMVM (100 nM). The data are presented as percent of remaining NADPH-oxidase activity in the presence of antagonists as compared to the responses from control cells. Quantification of data from 3 independent experiments are shown (mean ± SD, n=3). One-way ANOVA followed by Dunnett’s post-hoc test was used to calculate significance. **D)** Neutrophils were first stimulated with RE-04-001 (10 nM, arrow to the left) and then further challenged a second stimulation with WKYMVM (100 nM), fMLF (100 nM) or RE (10 nM) as indicated. **E)** Naïve or TNFα (37°C, 20 min) primed neutrophils were challenged with RE (10 nM). One representative trace out of three independent experiments is shown. **Inset:** Comparison between the peak O_2_^−^-responses released from naïve (black bars) and TNFα primed cells (grey bars) for either RE (10 nM), fMLF (100 nM) or WKYMVM (100 nM). Quantification of data from 3 independent experiments are shown (mean ± SD, n=3). Paired t-test was used to calculate the significance of TNFα priming effect. **F)** Comparison between the peak O_2_^−^ responses released by neutrophils, pre-treated with or without the Gαq inhibitor YM-254890 (200 nM) for five minutes before activation with RE (10 nM), fMLF (100 nM), WKYMVM (100 nM) or PAF (100 nM). The data are presented as percent of remaining NADPH-oxidase activity in the presence of YM-254890, compared to the responses from control cells, from 3 independent experiments (mean ± SD, n = 3). Paired t-test was used to calculate the significance of the effect of YM-254890.

The preference of RE for FPR1 over FPR2 in human neutrophils gained further support from receptor homologous desensitization experiments. Neutrophils first activated with RE were not only homologously desensitized (non-responsive) to a second stimulation with RE but were also refractory to stimulation with fMLF (Fig 3D). In contrast, these RE desensitized cells were still fully responsive to a second stimulation with the FPR2 agonist WKYMVM (Fig 3D). Taken together, these data show that RE is a very potent stimulus that activates the neutrophil NADPH-oxidase and this activation is achieved through signals specifically generated by FPR1.

It is well-known that the NADPH-oxidase activity triggered by FPR specific agonists is substantially increased in TNFα primed neutrophils [35, 45]. Accordingly, compared to the response induced in naïve neutrophils, also the amount of O_2_^−^ produced by TNFα primed cells were substantially increased with RE as the activating FPR agonist (Fig 3E). The increase due to priming with TNFα was of the same magnitude as that with fMLF and WKYMVM (Fig 3E inset). Finally, in agreement with the lack of inhibitory effect of the Gαq inhibitor on the FPR-mediated rise in [Ca^2+^]_i_ (Fig 2D), the O_2_^−^ production induced by RE and other FPR agonists (i.e., peptides fMLF and WKYMVM) was not inhibited by the Gαq inhibitor YM-254890 (Fig 3F). The inhibitory effects of the Gq inhibitor on PAF-induced NADPH-oxidase activity is shown for comparison (Fig 3F). Taken together, these data show that RE is a potent and full agonist selective for FPR1, and the agonist activates the neutrophil NADPH-oxidase independent of coupling to a Gαq containing G protein.

### 3.3 RE is less potent than fMLF to induce neutrophil migration

Based on the fact that the prototype FPR1 agonist fMLF and a large number of other earlier described FPR agonists are potently recruit migrating neutrophils, the FPRs are termed chemoattractant receptors [3, 4]. This generalization is, however, not completely valid based on the recent work with some FPR2 agonists such as lipidated peptides (pepducins and peptidomimetics) and the fomylated peptides belonging to the group of phenol soluble modulins (PSMα peptides). These agonists despite being potent activators in promoting superoxide release, they completely lack the ability to induce neutrophil chemotactic migration [30-32]. To determine the chemotactic activity of RE, we used the transwell chamber system in which neutrophils (placed in the upper chamber) were allowed to migrate through a filter that separates the agonist (placed in the bottom well in the chambers) from the cells. The FPR1 peptide agonist fMLF was used as positive control (Fig 4A). RE attracted neutrophils to a level similar to that by fMLF (Fig 4B), but comparably high concentrations were required to reach that level (Fig 4B), suggesting that signaling down-stream RE activated FPR1 is functional selective is in favor of oxidase over chemotaxis. Functional selective or biased signaling ratios were calculated to compare the functional selective profile of RE with that of fMLF; a ratio 0.1 and 5 was obtain for RE and fMLF, respectively, a value calculated a by a direct comparison of the respective EC_50_ value for activation of the NADPH-oxidase with that to recruit neutrophils chemotactically. Although the migration induced by a 50 nM concentration of RE reached the same level as that obtained with the optimal concentration of fMLF, the difference between the two agonist in the functional selective ratio values, clearly show that RE induced a functional selective response, being biased towards ROS production (Fig 3B) and away from chemotaxis (Fig 4B).

**Fig. 4.**
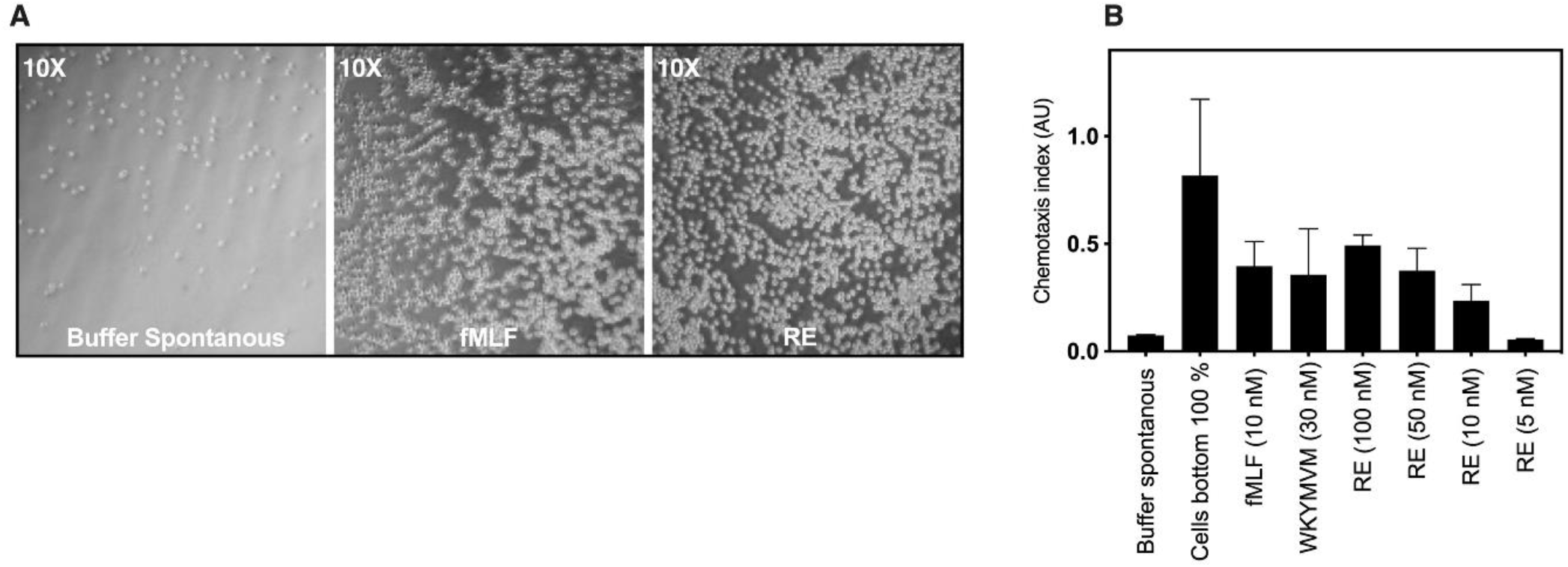
The small compound RE is a weaker chemoattractant than fMLF to induce neutrophil migration. Neutrophil migration was measured in a transwell chamber system where neutrophils were placed on the tope and agonists were placed in the bottom wells to allow migration during 90 min period at 37°C. **A**) Representative micrographs (10x magnification) of neutrophils recovered in the lower compartment after migration towards buffer control (spontaneous migration), fMLF (10 nM) or RE (50 nM). **B)** Migration of neutrophils towards fMLF (10 nM), WKYMVM (30 nM) or different concentrations of RE that was placed to the lower compartment. Neutrophils placed at the bottom chamber were regarded as 100% migration. The number of cells in the lower compartment was determined by analyzing the amount of MPO and shown as chemotaxis index from 3 independent experiments (mean + SD, n=3).

### 3.4 Biased neutrophil signaling by RE in favor of ERK1/2 phosphorylation over β-arrestin recruitment

The functional selectivity profile of RE in human neutrophils suggests that the agonist triggers a biased signal cascade downstream FPR1. In addition to a rise in [Ca^2+^]_i_, many FPR agonists trigger also ERK1/2 phosphorylation and recruitment of cytosolic β-arrestin to cytoplasmic parts of the activated receptors [46]. For many GPCRs the latter event is of importance for receptor desensitization and internalization as well as for the transduction of non-canonical signals of which activation of ERK1/2 may be one [47]. Phosphorylation of ERK1/2 in human neutrophils upon agonist stimulation was determined as previously described [32]. Similar to potent agonistic activity by RE in inducing a rise in [Ca^2+^]_i_, the agonist potently induced also ERK1/2 phosphorylation and the potency was comparable to, or slightly higher than that of fMLF (Fig 5A). The ability of RE to promote receptor-mediated recruitment of β-arrestin was studied in CHO cells overexpressing FPR1 [31]

**Fig. 5.**
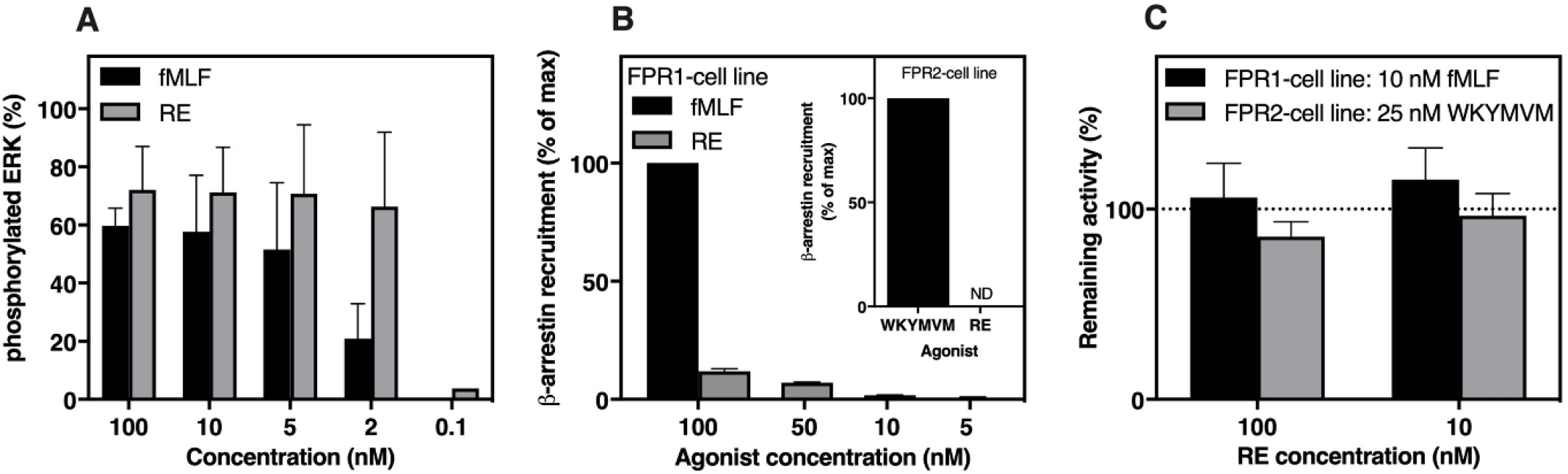
The small compound RE potently triggers ERK1/2 phosphorylation but poorly recruits β-arrestin. **A)** Phosphorylation of ERK1/2 (pERK1/2) was measured in neutrophil lysates using an MSD multispot assay system. Neutrophils were stimulated with different concentrations of fMLF or RE as indicated for 2 min followed by adding ice cold lysis buffer to stop phosphorylation process. The percent of phosphorylated ERK (% pERK) was calculated as: ([2xphospho-signal]/[phospho-signal + total signal]x100). The data are obtained from 2-3 independent experiments that were run with duplicates (mean + SD). **B)** β-arrestin recruitment monitored by PathHunter^®^ CHO cells over-expressing FPR1 or FPR2 together with β-arrestin 2. Response in FPR1 expressing CHO cells where stimulated with 100 nM of fMLF or different concentrations of RE as indicated. The data are presented as percent of the maximal response induced by a saturating concentration of fMLF (100 nM) for FPR1 expressing cells (mean + SD, n=3). **Inset:** FPR2 over-expressing CHO cells were stimulated with the FPR2 agonist WKYMVM (100 nM) or RE (100 nM). Data are presented as percent of the maximal response induced by 100 nM WKYMVM. **C**. FPR1 cells (black bars) and FPR2 cells (grey bars) were stimulated with fMLF (10 nM) and WKYMVM (25 nM), respectively, with or without the presence of RE (10 nM and 100 nM). The data are presented as percent of remaining β-arrestin recruitment in the presence of RE-04-001 as compared to the responses from control cells (mean + SD, n=3).

In contrast to the potent activity of RE in inducing a transient rise in [Ca^2+^]_i_ and ERK1/2 phosphorylation (Fig 2A, 5A), the amount of β-arrestin recruited by RE in FPR1 overexpressing cells was negligible in comparison to that induced by fMLF (Fig 5B). In agreement with the receptor specificity of RE, this agonist did not recruit any β-arrestin in FPR2 overexpressing cells; the FPR2 agonist WKYMVM was included as an FPR2 control for comparison (Fig 5B inset).

When comparing β-arrestin recruitment induced by the FPR1 agonist fMLF and RE, respectively, it is clear that whereas a full recruitment is achieved with a 10 nM concentration of fMLF, a very low level of β-arrestin recruitment (less that 20%) was obtained with much higher RE concentrations (Fig 5B). Despite the fact that RE is potent FPR1 agonist determined as a transient rise in [Ca^2+^]_i_, ERK1/2 phosphorylation and, activation of the NADPH-oxidase, RE did not block fMLF-induced β-arrestin recruitment (Fig 5C). As expected, RE lacked an effect also on FPR2 agonist WKYMVM-induced β-arrestin recruitment (Fig 5C). The functional selective profile of RE, away from chemotaxis and β-arrestin recruitment in comparison to its potent activity for ROS release and a transient rise in [Ca^2+^]_i_ as well as ERK1/2 phosphorylation, is basically in line with our earlier signaling profile of functional selective FPR2 agonists [30-32].

Taken together, these data clearly show that the FPR1 agonist RE displays not only a functional selectivity (NADPH-oxidase over chemotaxis) but also a strong signaling bias in favor of the signal giving rise to an increase in [Ca^2+^]_i_ and ERK1/2 phosphorylation over that resulting in β-arrestin recruitment.

### 3.5 RE promotes FPR1 to cross-talk with other neutrophil receptors

Following the response induced in neutrophils challenged with the FPR1 agonist fMLF or RE, the receptors/cells are transferred to a homologous desensitized state in which the cells are non-responsive to second agonist dose (Fig 3D). There is a known hierarchy between different neutrophil GPCRs, for example, FPR1 is ranked higher than the IL-8 receptors regarding both the NADPH-oxidase activation and neutrophil chemotaxis [48, 49]. In accordance with this, FPR1 homologous desensitization induced by fMLF and RE both resulted in a concomitant heterologous desensitization of the IL8 receptors, making these FPR1 desensitized cells non-responsive to IL-8 stimulation (Fig 6A).

**Fig. 6.**
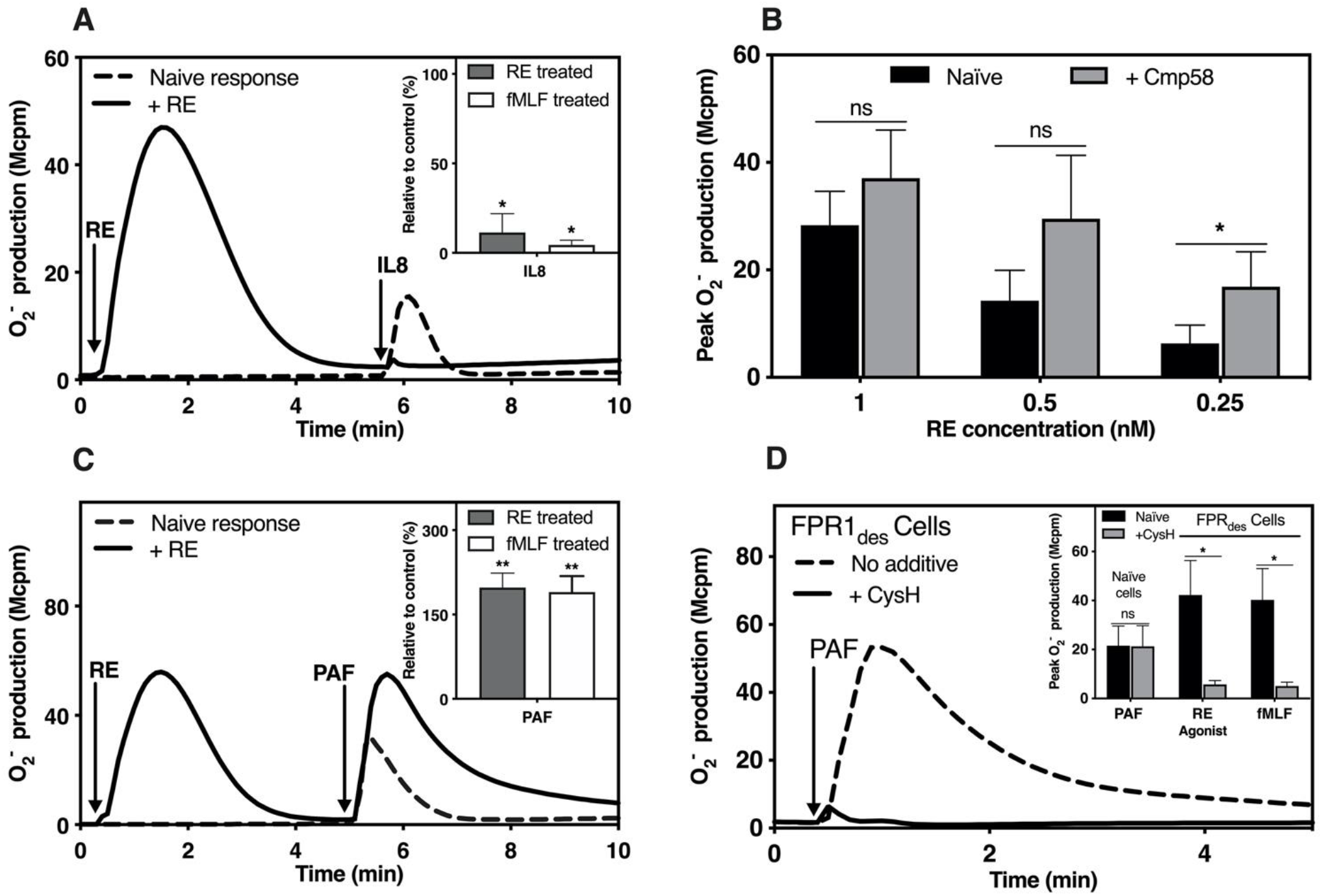
The small compound RE activated FPR1 modulates other GPCR-mediated neutrophil response. Receptor cross-talk was studied in the NADPH-oxidase activation assay by measuring the O_2_^−^ production. **A, C, D)** Abscissa, Time (min); ordinate, O_2_^−^ production in arbitrary Mcpm units. **A)** cross-talk between RE activated FPR1 and IL8. Neutrophils were activated with RE-04-001 (10 nM, indicated by the first arrow) when the response had declined, the cells received a second dose of IL8 (100 ng/ml; indicated by the second arrow). **Inset)** Quantification of the second IL8 response in RE or fMLF pre-activated cells from 3 independent experiments (mean + SD, n = 3). Paired t-test was used to calculate the difference from the naïve IL8 response. **B)** Different concentrations of RE were added to cells pre-treated with Cmp58 (1 μM), the FFAR2 allosteric modulator. Peak O_2_^−^ production from Cmp58 treated and naïve cells are shown from 3 independent experiments (mean + SD, n=3). Paired t-test was used to calculate the significance of the RE response obtained between Cmp58 treated and naïve cells. **C)** RE (10 nM, indicated by the first arrow) activated cells received a second stimulation with PAF (100 nM; indicated by the second arrow). One representative experiment out of three independent experiments is shown. **Inset)** The second PAF-response obtained from neutrophils pre-stimulated with either RE (10 nM) or fMLF (100 nM). Data are presented as % of control response from naïve neutrophils (mean + SD, n=4). Paired t-test was used to calculate the significance of the PAF response between FPR1 pre-activated and naïve cells. **D)** Neutrophils were desensitized with RE (10 nM) to obtain FPR1_des_ Cells before a second stimulation with PAF (100 nM). The FPR1 antagonist CysH was added just prior PAF stimulation (solid line) or cells received no addition before PAF stimulation (dashed line). Representative traces of O_2_^−^ production is shown from 4 independent experiments. **Inset)** Effect of CysH on the PAF response in naïve cells, cells desensitized with RE and fMLF (100 nM) were shown. Peak O_2_^−^ production induced by PAF from both naïve cells and FPRdes cells with or without CysH are shown (mean + SD, n = 4). Paired t-test was used to calculate the inhibitory effect of CysH.

Recent research suggests that the receptor cross-talk hierarchy is complex and not only desensitized receptors but also allosteric modulated GPCRs can communicate with other receptors [46, 50, 51]. A prominent example of such a cross-talk is that FPRs signaling can be positively regulated by free fatty acid receptor 2 (FFAR2) as illustrated by the fact that neutrophils with their FFARs allosterically modulated are primed when activated by low (normally non-activating) concentrations of FPR agonists [52, 53], and this is valid also for RE (Fig 6B); this RE induced response is inhibited not only to an FPR1 antagonist but also by an antagonist specific for FFAR2. This clearly shows that the response is achieved through receptor cross-talk between FPR1 and FFAR2.

Opposite to the heterologous inhibitory effect of RE on the IL-8 response (Fig 6A), a substantially enhanced PAF response was induced in FPR1-desensitized neutrophils compared to the naïve PAF response (Fig 6C) and no difference was observed in cells when desensitized by fMLF or RE (Fig 6C inset). The involvement of FPR1 is this response is evident from the fact that the second PAF response in RE desensitized cells, is sensitive to the FPR1 antagonist cyclosporine H when added just prior to PAF stimulation (Fig 6D). This is in line with the earlier data showing that PAF/PAFR is able to transduce a not yet known signal leading to a reactivation of neutrophils with desensitized FPRs [54].

In summary, we show that the novel FPR1 agonist RE, despite its biased signaling feature, similar to fMLF places FPR1 in the same position in the neutrophil receptor hierarchy and allow receptor cross-talk with other GPCRs to either suppress or amplify the neutrophil response.

### 3.6 The termination of the RE induced activation of the NADPH-oxidase is regulated primarily by the actin cytoskeleton rather than by β-arrestin

Despite the fact that β-arrestin plays an important role in receptor desensitization for many GPCRs, we and others have demonstrated that the actin cytoskeleton, rather than the recruited β-arrestin, constitutes the basis for FPR desensitization and termination of signals that activate the ROS producing oxidase [46, 55, 56]. This notion gains further support from the fact that FPR1 is homologously desensitized also by RE, despite the fact that no β-arrestin is recruited by this agonist. In addition, in neutrophils pre-treated with the actin cytoskeleton disrupting agent latrunculin A, RE induced activation resulted in a 4-fold higher superoxide production in comparison to that produced by naive (untreated) cells (Fig 7A). Further, RE activated neutrophils transferred to a non-signaling desensitized state, were resensitized/reactivated and produce ROS when the actin cytoskeleton was disrupted through the addition of latrunculin A (Fig 7B). These data, obtained with RE as activating agonist, are in agreement with the pattern when fMLF was used as FPR1 agonist to activate-desensitize-resensitize/reactivate neutrophils (Fig 7A, B). Taken together, we show that RE-induced FPR1 desensitization in neutrophils, occurs primarily through the involvement of an intact actin cytoskeleton.

**Fig. 7.**
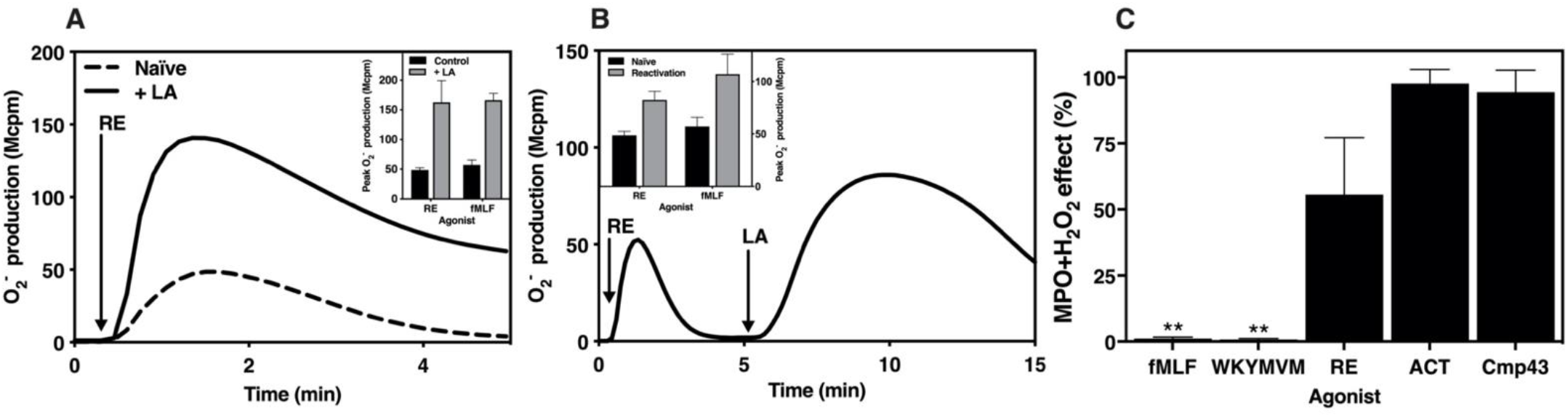
Modulation of the small compound RE activity by Latrunculin A and the MPO/H_2_O_2_ system. The NADPH-oxidase activity induced in neutrophils was determined Abscissa, Time (min); ordinate, O_2_^−^ production, arbitrary Mcpm units). **A)**. Naïve neutrophils and neutrophils incubated (five minutes) with the actin cytoskeleton-disrupting drug latrunculin A (LA; 25 ng/mL) were activated with RE (10 nM). One representative experiment of three independent experiments is shown. **Inset:** The peak NADPH-oxidase activities induced in naïve or LA treated neutrophils by RE (10 nM) or the FPR1 agonists fMLF (100 nM) were determined (mean + SD, n=3). **B)**. Neutrophils activated by RE (10 nM, addition indicated by the arrow to the left) were reactivated with LA (25 ng/mL, addition indicated by arrow to the right). One representative experiment of three independent experiments is shown. **Inset:** The peak NADPH-oxidase activity in naïve neutrophils induced by RE (100 nM), or the FPR1 agonist fMLF (100 nM) respectively were determined and compared with the peak NADPH-oxidase activity induced after reactivation with LA (mean + SD, n=3). **C)**. Sensitivity of the agonist towards MPO-H_2_O_2_ system induced oxidization. FPR agonists were exposed to MPO (1 μg/ml) + H_2_O_2_ (10 μM) and the remaining activity was measured by the degree of agonists to trigger ROS production from neutrophils. Final concentrations of agonist in the oxidase assay was fMLF (100 nM); WKYMVM (100 nM); RE (12.5 nM); Act-389949 (ACT, 12.5 nM) or Cmp43 (250 nM). The control agonists received no MPO-H_2_O_2_. The data are presented as percent of activity for each agonist treated with or without MPO-H_2_O_2_ from 3 independent experiments. Paired t-test was used to calculate the activity difference for each individual agonist treated with or without MPO-H_2_O_2_ (mean + SD, n = 3).

### 3.7 RE is resistant to oxidation by the MPO-H_2_O_2_-halide system

Processing of NADPH-oxidase-derived hydrogen peroxide (H_2_O_2_) by myeloperoxidase (MPO), a neutrophil protein stored in the azurophil granules, results in a generation of highly reactive oxidants that regulate many biological processes in addition to bacterial killing [15, 38, 57]. In line with this, the MPO-H_2_O_2_-halide system inactivates the peptide agonists fMLF and WKYMVM ([35, 58]; Fig 7C), as evident from the inability of the peptides to trigger ROS release from neutrophils. The small compound agonists Cmp43 and Act-389949 resisted completely the MPO-H_2_O_2_-halide radical system (([35, 59]); Fig 7C). Compared to the peptide FPR1 agonist fMLF, RE was fairly resistant to the MPO-H_2_O_2_-halide radical system (Fig 7C). Taken together, these data show that the RE resists inactivation induced by the MPO-H_2_O_2_-halide system.

## 4. Discussion

In this study, we show that RE, a small molecule that, by binding to FPR1, activates the neutrophil superoxide generation NADPH oxidase, and it is also shown that FPR1 selective recognizes this agonist. In-depth characterization of this FPR1 agonist reveal that there are large similarities regarding activation profiles between RE and the prototype peptide agonist fMLF, with the exception that the ability of RE to recruit neutrophils chemotactically is reduced, and this functional selectivity was associated with a weaker ability of to recruit β-arrestin. Despite the rapid progress in the identification of potent agonists for both FPR1 and FPR2, very few FPR1 selective agonists that display biased signaling and functional selective properties in human neutrophils have been described. Only a few FPR agonists have been progressed into clinical development, and at present, one FPR1/2 dual agonist Compound 17b has been reported to exert anti-inflammatory effects and protect mice from myocardial infarction injury [60], whereas another compound (BMS-986235) has developed into clinical phase I studies by Bristol Meyers-Squibb as a selective FPR2 agonist for prevention of heart failure [61]. Yet another FPR2 selective agonist Act-389949 entered a clinical phase I study but the data obtained show that the receptors exposed on the surface of blood neutrophils were rapidly lost but the precise mechanism for this was not described [62]. It is clear that better understanding of the basic biology and of the mechanisms that regulate FPRs is highly desirable as both the precise roles of FPR1 and FPR2, the effects of biased agonist as well as of receptor desensitization and the intracellular signals generated (including recruitment of β-arrestin) will be of importance for downstream cellular response and therapeutic effects.

Peptides with a formylated methionine in their N-terminus, a hallmark of protein/peptide synthesized by bacteria and mitochondria, are recognized by the innate immune system through high affinity binding of formylated peptides to FPR1 and/or FPR2, receptors expressed primarily in myeloid cells such as granulocytes and monocytes/macrophages [3, 4, 46]. Following early work showing that formylated peptides are high affinity FPR ligands, FPR1 as well as the closely related FPR2 have been shown to be promiscuous and recognize also a large number of compounds lacking the formylated methionine. Our identification of RE as a novel FPR1 agonist is in line with the promiscuous ligand binding feature for FPR1, the first one of the GPCRs expressed in neutrophils to being cloned, and much our knowledge about this receptor has been obtained with the high affinity bacterial-derived peptide fMLF, a commonly used research tool [4]. Shortly after the cloning and deorphanization of FPR2, extensive research highlighted a promiscuous ligand binding profile also for this FPR, i.e., both receptors bind with high affinity, formylated peptides of bacterial as well as host cell origin, and numerous non-formylated molecules belonging to different structural classes [9, 46]. Some of these ligands have overlapping binding profiles but other are selectively recognized by one or the other of the two FPRs. The precise structural requirements for ligand recognition by the FPRs are still poorly understood, but interestingly, two very recent structure biology studies have revealed the crystal structure of FPR2 in its active conformation in complex with the high affinity peptide agonist WKYMVm [63, 64]. Future molecular docking of FPR1 selective agonists using an FPR2-based model of FPR1, may define the mechanistic insights into FPR1 selective recognition of such compounds. These types of studies may also reveal differences in conformational and binding modes between fMLF and RE, agonists that transduce distinct signaling pathways and trigger different cellular responses (see discussion below).

It is generally accepted that activation by receptor specific agonists of chemoattractant GPCRs such as the FPRs, regulates the recruitment of neutrophils from the blood stream to inflammatory sites in infected/damaged tissues and the receptor down-stream signals generated induce also a release/secretion from these cells of proteolytic enzymes and ROS [3]. We show that similar to the FPR1 agonist fMLF, RE acts as a full agonist for activation of the ROS generating NADPH-oxidase, and the level of ROS production is largely amplified/primed in cells pre-treated with TNFα. The precise molecular background to the TNFα primed response is not known but it may be the result of an increased exposure of membrane receptors mobilized from stores in the secretory granules. Our earlier studies have demonstrated that such secretory organelles containing CD11b, FPR1and, FPR2 are mobilized to the neutrophil surface by priming agents such as TNFα and LPS [65-67]. Considering the high levels of TNFα in a number of inflammatory diseases such as rheumatoid arthritis, suggests that the mechanism underlying the neutrophil priming process and its consequences both *in vitro* and *in vivo* may offer new opportunities for therapeutic intervention in pathological settings [68].

Despite the robust release of ROS induced by low nM concentrations of RE, much higher concentrations were needed to induce neutrophil chemotaxis. Thus, RE clearly reveals a functional selective neutrophil response. This biased signaling profile of this functional selective FPR1 agonist is in large the same as that of earlier described for functional selective FPR2 agonist such as bacterial-derived PSMα peptides, pepducins and lipidated peptidomimetics. These FPR2 agonists activate the neutrophil ROS producing NADPH-oxidase but lack the ability to recruit neutrophils chemotactically [30-32]. Taken together these data suggest that both FPR1 and FPR2 at the molecular level, can be stabilized in conformations that open for one signaling pathway but not for another, a signaling bias that gives rise to a functional selective response, a signaling profile/functional outcome determined by the binding mode of the agonist. This suggestion is also supported by data obtained with variants of the prototype peptide agonist fMLF, that have been shown to trigger chemotaxis but be unable to activate the ROS generating neutrophil NADPH-oxidase [33]. Future structural studies of FPR1 in complex with different agonists should provide molecular insights into the ligand-directed FPR1 activation mechanism.

The FPR1 signaling scheme for RE includes the signals that induce a transient rise in [Ca^2+^]_i_, one of the very early events in GPCR signaling, and based on the activity induced by RE, it is clear that this agonist is more potent than the prototype FPR1 agonist fMLF. The increase in [Ca^2+^]_i_ is not reduced by a Gαq-inhibitor, and this is in line with earlier studies that have identified the βγ part of a Gαi containing G protein downstream of FPR1, to be the link between the receptor an activation of the PLC-PIP_2_-IP_3_-Ca^2+^ pathway [4, 44]. Similar signaling profiles of fMLF and RE are obtained also for the receptor down-stream signal leading to an activation of ERK1/2 phosphorylation; i.e., it is clear that RE is a more potent agonist than the prototype peptide agonist fMLF. Despite this, we noticed an obvious difference between the two agonists with respect to their ability to activate FPR1 for β-arrestin recruitment, demonstrating a biased signaling profile downstream of FPR1 when activated with RE.

The biased signaling concept is now firmly established in GPCR biology [27-29]. Clearly, this concept is valid also for FPR1; in contrast to the prototype FPR1 agonist, RE has a biased signaling profile as illustrated by the fact that despite ability to potently activate ERK1/2 phosphorylation and induce a rise in [Ca^2+^]_i_, a very low level of β-arrestin recruitment was induced by RE. Similar to RE, several FPR2 agonists have earlier been shown to transduce a biased signaling feature in neutrophils [30-32]. It is interesting to note that similar to RE, the biased signaling FPR2 agonists that lack ability to recruit β-arrestin and are also poor neutrophil chemoattractants [30-32], suggesting a role for β-arrestin in regulating both FPR1- and FPR2-mediated directional cell migration. The non-peptide compound termed Quin-C1 [69] has also been shown to be a biased signaling FPR2 agonist, but with the reversed functional selectivity; it lacks the ability to trigger superoxide release, while being able to induce neutrophil chemotaxis [69]. The precise mechanism of this type of biased signaling down-stream of a receptor occupied by different ligands is not clear at present, but the molecular basis for this phenomenon has been suggested to be due to the formation of different receptor subconformations induced by different agonist, which in turn transduce different strength of signaling pathways and cellular responses. It should also be noticed that β-arrestin has been suggested to regulate receptor desensitization and internalization as well as to initiate non-canonical signaling including ERK1/2 phosphorylation [47, 70]. Our data showing that RE at concentrations that are unable to recruit β-arrestin is despite this a potent trigger of the ERK1/2 phosphorylation pathway, suggesting that FPR1-mediated ERK1/2 phosphorylation is not regulated by β-arrestin.

Although there are differences in the activation/signaling profiles between RE and the prototype peptide FPR1 agonist fMLF, our data also demonstrate that there are similarities; RE similar to fMLF, interplay with other neutrophil GPCRs; this is achieved through different GPCR cross-talk mechanisms, complex phenomena with mechanisms not yet understood (see a recent review, [46]). Nevertheless, the biological relevance of receptor cross-talk is obvious when neutrophils facing multiple ligands that have affinity for different receptors during migration and activation process. The outcome of neutrophil activation is thus dependent on the co-operation of multiple ligands at the receptor signaling level. This co-operation is evident from our data demonstrating that RE can inhibit IL-8 but prime the PAF response. When it comes to the cross-talk between FPR1 and FFAR2, low concentrations of RE could be primed by allosterically modulated FFAR2. The fact that the primed PAF response is sensitive to an FPR1 antagonist further supporting the cross-talk mechanism relies on a reactivation of desensitized FPR1 [54, 71].

In summary, we have identified and characterized a small compound as a potent FPR1 selective agonist, and we provide some unique feature of RE in triggering biased FPR1 signaling and neutrophil functional selectivity, activation characteristics that differ from the most commonly used FPR1 peptide agonist fMLF. The information provided about the basic characteristics of RE should be of value for further optimization processes and mechanistic studies both *in vivo* and *in vitro* and the knowledge obtained would shed more light on the complex biology of FPR1 in health and in different disease conditions.

## Acknowledgements

This work was supported by the Swedish Research Council, the Knut and Alice Wallenberg foundation, the Swedish government under the ALF-agreement, the IngaBritt and Arne Lundberg research foundation, Eurostars project rosa GBS-E 10082, and Swedish Governmental Agency for Innovation Systems (project number 2016-01010). This research has received funding from the [European Union’s Horizon 2020 research and innovation program project NEUTROCURE under grant agreement No 861878.

## Authorship contributions

SL and HF performed the experiments and analyzed the data with the input from the co-authors. RH, PO, CD and HF designed the study and analyzed the data. HF and CD wrote the manuscript and all authors commented on before approving the final version.

